# Activin A correlates with the worst outcomes in COVID-19 patients, and can be induced by cytokines via the IKK/NF-kappa B pathway

**DOI:** 10.1101/2021.02.04.429815

**Authors:** Megan McAleavy, Qian Zhang, Jianing Xu, Li Pan, Matthew Wakai, Peter J. Ehmann, Matthew F. Wipperman, Tea Shavlakadze, Sara C. Hamon, Anita Boyapati, Lori G. Morton, Christos A. Kyratsous, David J. Glass

## Abstract

A fraction of COVID-19 patients develop the most severe form, characterized by Acute Respiratory Disease Syndrome (ARDS). The molecular mechanisms causing COVID-19-induced ARDS have yet to be defined, though many studies have documented an increase in cytokines known as a “cytokine storm.” Here, we demonstrate that cytokines that activate the NF-kappaB pathway can induce Activin A and its downstream marker, FLRG. In hospitalized COVID-19 patients elevated Activin A/FLRG at baseline were predictive of the most severe longitudinal outcomes of COVID-19, including the need for mechanical ventilation, lack of clinical improvement and all-cause mortality. Patients with Activin A/FLRG above the sample median were 2.6/2.9 times more likely to die, relative to patients with levels below the sample median, respectively. The study indicates high levels of Activin A and FLRG put patients at risk of ARDS, and blockade of Activin A may be beneficial in treating COVID-19 patients experiencing ARDS.

## Introduction

In the setting of infection by the SARS-Cov2 virus, it was reported quite early that hospitalized and ICU patients were producing a “cytokine storm” (Huang et al., 2020), including the cytokines IL-1 and TNF*α*. Clinical studies have demonstrated that blockade of cytokine signaling and steroid treatment are beneficial in improving outcomes in patients, however further elucidation of downstream signaling pathways contributing to clinical sequelae is important to benefit patients suffering the worst symptoms of Covid-19.

We had previously studied IL-1 and TNF*α* in the setting of skeletal muscle cachexia, where these cytokines have been shown to induce skeletal muscle atrophy (Egerman and Glass, 2014; Trendelenburg et al., 2012). In one of our prior studies, we learned that IL-1 and TNF*α* could induce the production of Activin A in skeletal muscle, and that the Activin A itself induced skeletal muscle atrophy. We felt this was relevant to COVID-19, because it had been reported separately, back in 2012, that patients who had Acute Respiratory Disease Syndrome (ARDS), had high levels of Activin A in their Bronchial Alveolar Lavage fluid (Apostolou et al., 2012), and, in a preclinical model, this same group found Activin A to be sufficient to induce a phenotype reminiscent of ARDS when over-expressed in the trachea via an Adenovirus (Apostolou et al., 2012). A separate group followed up in 2019, on a distinct ARDS population, and were able to show that Activin A and its downstream pathway marker, FLRG, were upregulated in human serum (Kim et al., 2019).

In addition, the most severe symptoms associated with Covid-19 seem to be age-related; older patients and those with particular co-morbidities, like COPD, are more likely to experience ARDS and are at higher risk for mortality from the virus (Fang et al., 2020; Wang et al., 2020). It’s therefore of interest to determine molecular mechanisms which are themselves age-perturbed, which might help to explain this correlation of aging with Covid19-induced mortality.

For these reasons, we were interested to study sera from COVID-19 patients, to determine if they too had elevated levels of Activin A, and evidence of pathway elevation, correlated to FLRG levels. In addition, another marker previously associated with ARDS, PAI-1, was also evaluated as it is one of the parameters confirmed in the ARMA and ALVEOLI trials associated with ARDS mortality (Bos et al., 2017; Calfee et al., 2015). We further sought to determine if the levels of Activin A, FLRG and PAI-correlated to important disease markers of COVID-19, such as disease severity, the requirement for supplemental oxygen, other signs of ARDS, and mortality. On a mechanistic level, we were then interested to see if other COVID-19 relevant cell types, including bronchial and pulmonary smooth muscle, similarly responded to inflammatory cytokines induced by the Cytokine storm, to produce Activin A, and, if so, by which signaling pathway.

We had performed a clinical trial on COVID-19 patients using a Regeneron anti-IL-6R antibody (sarilumab) (https://clinicaltrials.gov/ct2/show/NCT04315298). We evaluated sera from these patients after randomization and prior to therapy, to determine baseline Activin A, FLRG and PAI-1 levels, and correlated these to baseline clinical and laboratory variables and important disease outcomes.

## Results

### Activin A, FLRG, and PAI-1 are elevated in critical patients relative to severe patients or healthy controls

COVID-19 presents a full spectrum of disease severity, from asymptomatic, mild cold-like symptoms, more disabling but ambulatory illness, to more severe illness requiring degrees of hospitalization and ICU Care, including increasing levels of oxygen support or ventilation. In order to evaluate the relationship between Activin pathway engagement and stages of severe disease progression, we examined the levels of Activin A and its pathway marker FLRG in sera from COVID-19 hospitalized pneumonia patients with varying disease severity. Baseline samples were collected from individuals hospitalized requiring low to high supplemental oxygen who were receiving standard of care and supportive therapies due to Sarilumab trial enrollment early in the pandemic at US centers. These were collected following randomization and prior to treatment. To contextualize levels in COVID-19 patients, age-matched sera from healthy controls were also evaluated.

In a prior study, it was shown that Activin A can be found in the Bronchial Alveolar Lavage fluid and in the sera of other, non COVID-19, ARDS patients (Apostolou et al., 2012). In a preclinical model, it was further shown that, when over-expressed, Activin A was sufficient to phenocopy ARDS (Apostolou et al., 2012). Therefore, we sought to determine if Activin A and its pathway marker FLRG, are elevated in severe and ICU COVID-19 patients in comparison to healthy controls. SARS-CoV2-infected sera was obtained from subjects in a clinical study, where age-matched sera samples from healthy controls were purchased. Additionally, PAI-1 was studied because it is involved in the coagulation pathway (Ahmed et al., 2020), so it seemed reasonable to use it as an additional queried biomarker of interest in COVID due to its multiple roles in inflammation, coagulation, as well as an Activin-response protein, to determine if there was anything unique about the Activin A/FLRG pathway.

### Baseline demographics of biomarker study population

Demographic and health characteristics of the COVID-19 patients studied are shown in Table 1. From the overall study, which enrolled 1946 patients from 62 sites, the present analysis includes a random subset of 312 COVID-19 patients from 49 sites in addition to 114 age-matched control subjects. Patients in the critical disease stratum had higher percentage of patients admitted to the ICU and receiving steroids or vasopressors compared to patients in the severe stratum (Table 1).

**Table 1.**
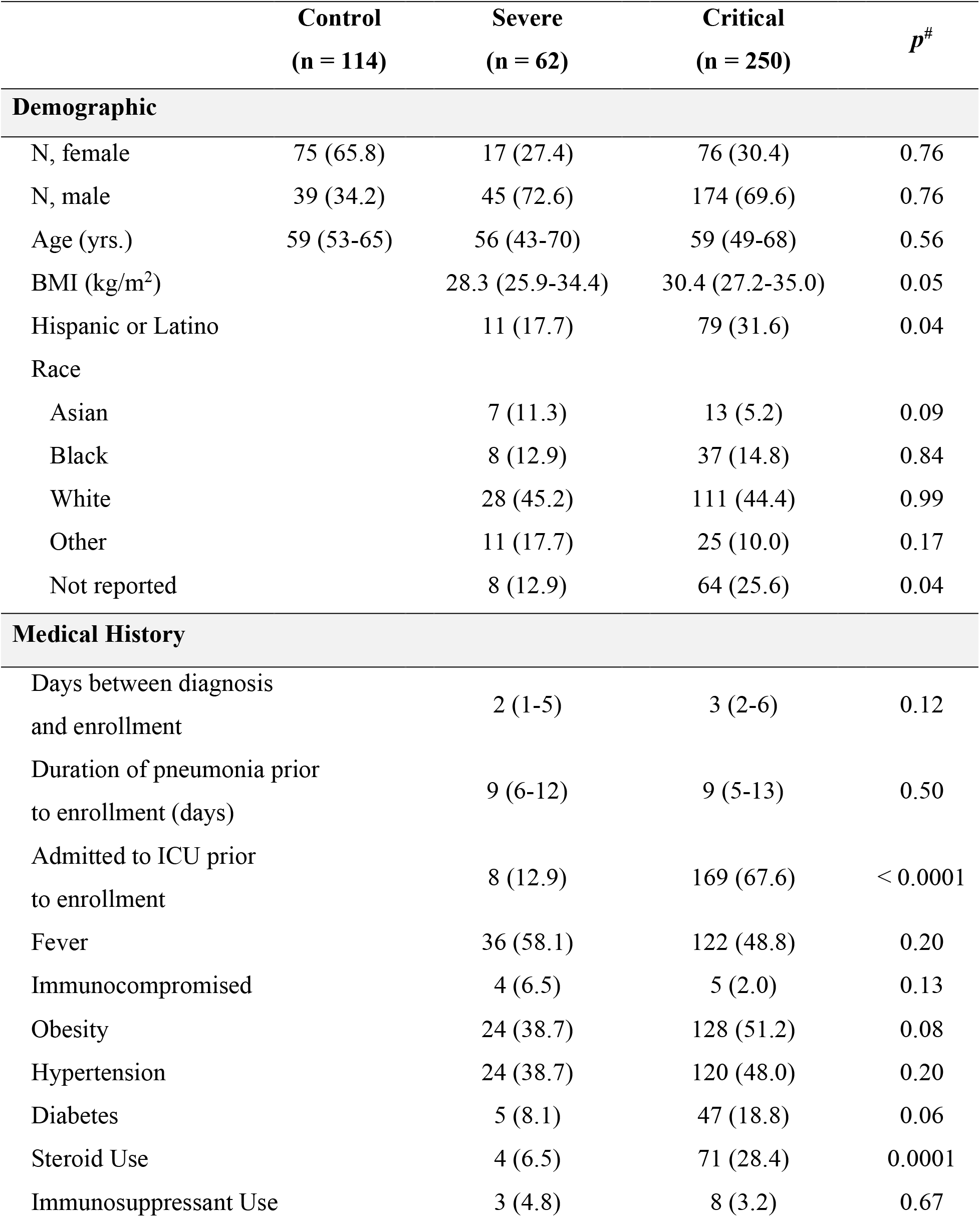
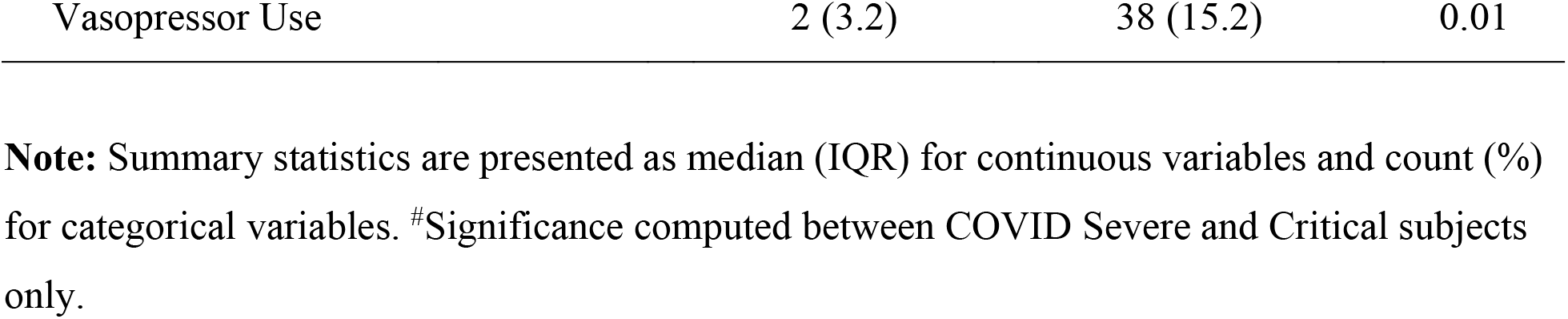
**Demographic information and medical history grouped by disease severity**.

### Activin A, FLRG, PAI-1 not affected by sarilumab treatment relative to placebo

Although the clinical data analysis focused on pre-treatment sera analysis, we tested for potential pharmacodynamic effects of sarilumab on Activin A, FLRG, and PAI-1 that may confound prediction of longitudinal outcomes. Due to sample limitations, only 182 patients’ (n=95 placebo, n=87 sarilumab 400mg) baseline samples were analyzed along with matching samples at Day 4 (N=176; n=92 placebo, n=84 sarilumab 400mg) and/or Day 7 (N=143; n=59 placebo, n=84 sarilumab 400mg). These data are only from patients in the Phase 3 study (Critical disease severity). For Activin A, FLRG and PAI-1, there was no significant difference between treatment arms at baseline and change from baseline at Day 4 or Day 7 (data not shown).

### Activin A and FLRG are elevated in patients with greater disease severity

Differences were observed between three categories of disease severity (control subjects, severe COVID-19 patients, critical COVID-19 patients) for Activin A, FLRG, and PAI-1 (Figure 1; *p* < 0.0001). Follow-up pairwise testing revealed significantly elevated Activin A in critical (ICU) COVID-19 patients, as compared to severely affected non-ICU COVID-19 patients and control subjects (*p* < 0.05/3); severe non-ICU COVID-19 patients and control subjects did not differ in Activin A (*p* > 0.05/3), indicating that Activin A is a biomarker for the Critical, ICU-bound patients. For FLRG, levels were significantly elevated with increased severity of disease; control, Severe, and Critical (*p* < 0.05/3), indicating that this Activin A pathway marker can also distinguish between the distinct disease categories. In contrast, for PAI-1, levels were not statistically different between the two COVID-19 strata: Severe and Critical (*p* > 0.05/3). However, both were significantly elevated compared to control subjects (*p* < 0.05/3). These data demonstrate that Activin A and its pathway marker FLRG are upregulated in more severe settings of COVID-19, where patients require treatment in the ICU. In contrast, PAI-1 was elevated in both COVID-19 settings and did not distinguish between the severe and critical state.

**Figure 1:**
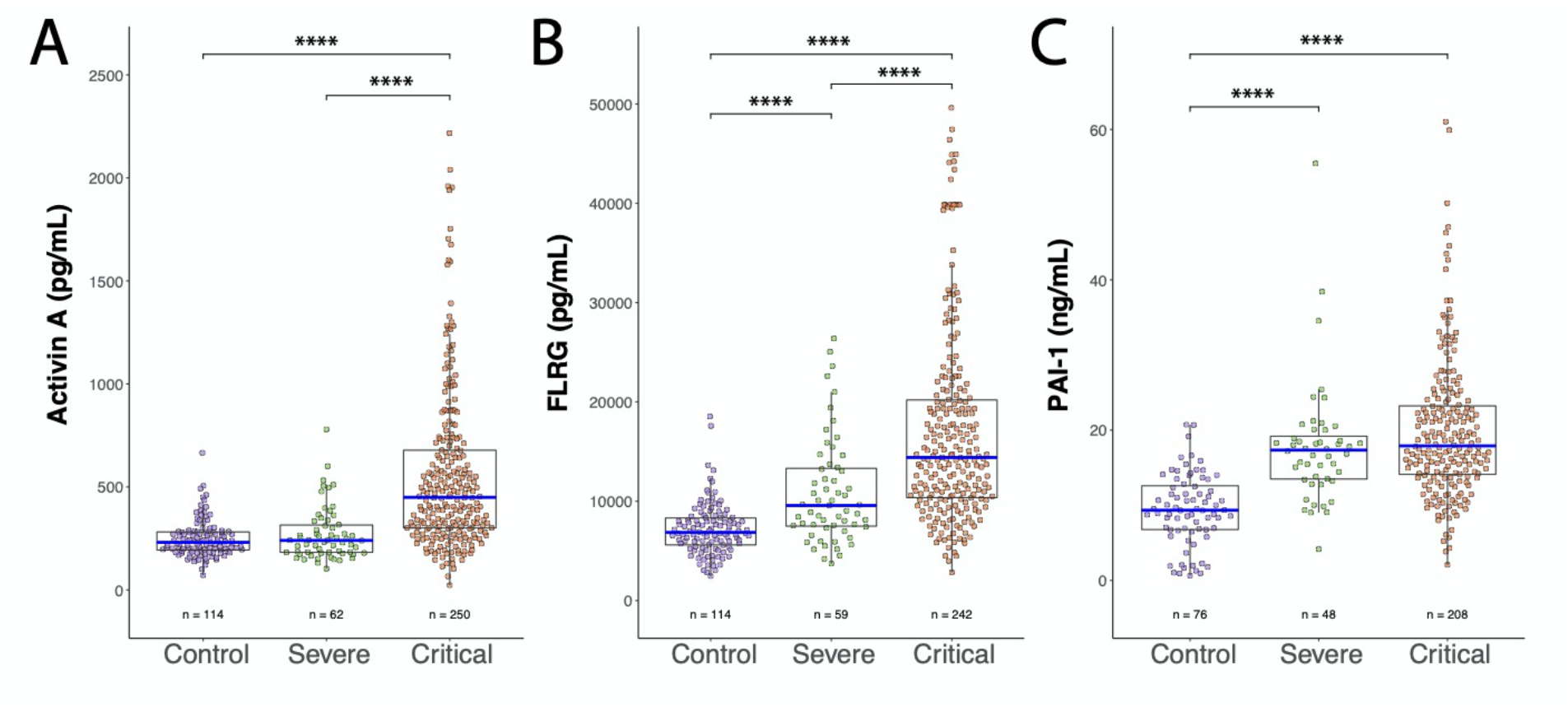
Activin A, FLRG, and PAI-1 levels vs disease severity in COVID-19 patients and in non-Covid 19 controls. **A**. Activin A (pg/mL) levels plotted for control, severe COVID-19, and critical COVID-19 subjects. Significant differences between control and critical COVID-19, and severe and critical COVID-19 were found. **B**. FLRG (pg/mL) levels plotted as in A. All groups were significantly different from each other, with FLRG levels increasing with disease severity. **C**. PAI-1 (ng/mL) levels plotted as in A. Significant differences were found between control and severe COVID-19, and control and critical COVID-19. Number of subjects tested in each group (n) is indicated under respective plots. **** *p* < 0.0001.

### Higher levels of Activin A & FLRG are associated with increased risk of death and greater oxygen requirements at baseline

Since Activin A and FLRG levels were most highly elevated in ICU patients (Fig. 1) we were further interested in examining the relationship between longitudinal outcomes with baseline Activin A, FLRG, and PAI-1 in COVID-19 patients (Fig. 2 A-C). Baseline Activin A was lower in patients who survived than in patients who died and was a significant predictor of all-cause mortality (OR = 1.54 (1.22, 1.98), *p* = 0.0004). The same trend was observed for FLRG (OR = 1.69 (1.32, 2.18), *p* < 0.0001), which was also lower in patients who survived than in patients who died, consistent with Activin A and FLRG as being part of the same pathway - FLRG is induced by Activin A activation of Smad2/3 (Razanajaona et al., 2007). In contrast, baseline PAI-1 was not predictive of mortality (OR = 1.19 (0.92, 1.55), *p* = 0.18).

**Figure 2:**
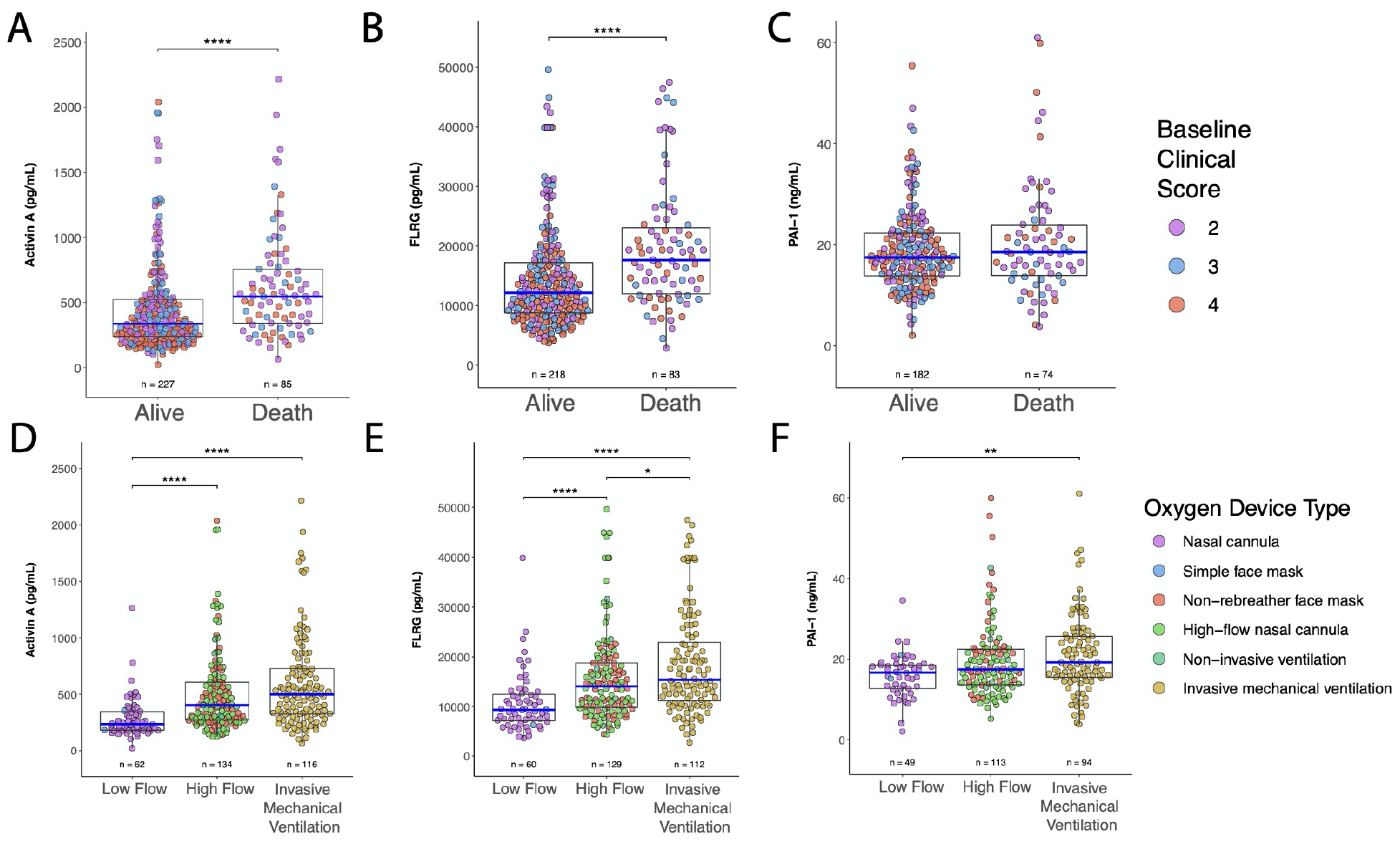
Activin A and FLRG levels associate with mortality outcomes and supplemental oxygenation requirements at baseline. **A**. Activin A levels (pg/mL) at baseline between alive vs dead subjects. **B**. FLRG levels (pg/mL) at baseline between alive vs dead subjects. **C**. PAI-1 levels (ng/mL) at baseline between alive vs dead subjects. **D**. Activin A levels (pg/mL) vs baseline supplemental oxygen requirements. **E**. FLRG (pg/mL) levels vs baseline supplemental oxygen requirements. **F**. PAI-1 (ng/mL) levels vs baseline supplemental oxygen requirements. Number of subjects analyzed in each group (n) is indicated under respective plots. * *p* < 0.05, ** *p* < 0.01, **** *p* < 0.0001.

Oxygenation status at baseline was stratified into three categories based on oxygen device type: Low Flow, High Flow, and Invasive Mechanical Ventilation (IMV). Individual subject data are plotted in Figure 2 (D-F). Differences between these three groups were observed for Activin A (*p* < 0.0001), FLRG (*p* < 0.0001), and PAI-1 (*p* = 0.0004). Activin A was lowest for patients on Low Flow, higher for patients on High Flow, and highest for patients on IMV. However, Activin A did not significantly differ between patients on High Flow and IMV (*p* > 0.05/3). The same trend was observed for FLRG and all pairwise comparisons were significant (*p* < 0.05/3). PAI-1 was also elevated with increased oxygen requirements. However, PAI-1 levels were only significantly different between patients on Low Flow and IMV (*p* < 0.05/3). These data indicate that Activin A and its pathway marker, FLRG, correlate with need for higher oxygen levels, but PAI-1 levels, which is generally elevated in COVID-19 patients relative to healthy subjects, are not linked with the need for greater oxygen.

In addition to Activin A and FLRG, disease progression and clinical outcomes also were related to supplemental oxygen requirements upon study enrollment (Table 2). Patients requiring greater supplemental oxygen pre-treatment experienced more days with fever, tachypnea, hypoxemia, and supplemental oxygen. Furthermore, elevated rates of mortality, lower rates of clinical improvement, and lower rates of hospital discharge were observed in patients requiring greater oxygen requirements.

**Table 2.** **Activin pathway laboratory resultsand clinical outcomes grouped by baseline supplemental oxygen requirements**.

|  | **Low Flow**<br>**(n = 62)** | **High Flow**<br>**(n = 134)** | **Invasive**<br>**Mechanical**<br>**Ventilation**<br>**(n = 116)** | ***P*** |
| --- | --- | --- | --- | --- |
| **Activin Pathway** |  |  |  |  |
| Activin A (pg/mL) | 236.5<br>(182.2-346.1) | 403.7<br>(271.7-607.5) | 499.6<br>(324.9-726.9) | < 0.0001 |
| FLRG (pg/mL) | 9399<br>(7233-12497) | 14117<br>(9889-18800) | 15424<br>(11239-22936) | < 0.0001 |
| PAI-1 (ng/mL) | 16.7<br>(12.7-18.6) | 17.5<br>(13.7-22.5) | 19.2<br>(15.5-25.6) | 0.0004 |
| **Clinical Outcomes** |  |  |  |  |
| All-cause mortality | 8 (12.9) | 33 (24.6) | 45 (38.8) | 0.0003 |
| Clinical score<br>improvement ($\geq 1$pt) | 51 (82.2) | 88 (65.7) | 58 (50.0) | < 0.0001 |
| Improvement in<br>oxygenation | 44 (71.0) | 92 (68.7) | 65 (56.0) | 0.07 |
| Discharge | 50 (80.6) | 83 (61.9) | 46 (39.7) | < 0.0001 |
| Number of days with<br>fever <sup>#</sup> | 1 (0-1) | 1 (0-2) | 3 (1-7) | < 0.0001 |
| Number of days with<br>tachypnea <sup>#</sup> | 0 (0-1) | 0 (0-2) | 5 (2-12) | 0.0009 |
| Number of days with<br>hypoxemia <sup>#</sup> | 7 (4-10) | 12 (8-20) | 19 (13-32) | < 0.0001 |
| Number of days on<br>supplemental O<sub>2</sub> <sup>#</sup> | 7 (4-10) | 12 (8-19) | 19 (10-30) | < 0.0001 |

**Note:** Summary statistics are presented as median (IQR) for continuous variables and count (%) for categorical variables. <sup>#</sup> variables assessed only in survivors.

### Lower Activin A and FLRG are associated with Lower rates of Mortality and 1-point improvement in Clinical score

We next assessed the effect of baseline Activin A, FLRG, and PAI-1 on clinical endpoints, including all-cause mortality and clinical score improvement (*≥*1 point), using Fine-Gray subdistribution hazard models. Patients were median split into two groups (Low, High) for each analyte (Activin A = 388.4 pg/mL, FLRG = 13531 pg/mL, PAI-1 = 17.7 ng/mL). Cumulative incidence curves for both endpoints are displayed in Figure 3 and subdistribution hazard ratios (sHR) are shown in Table 3.

**Figure 3:**
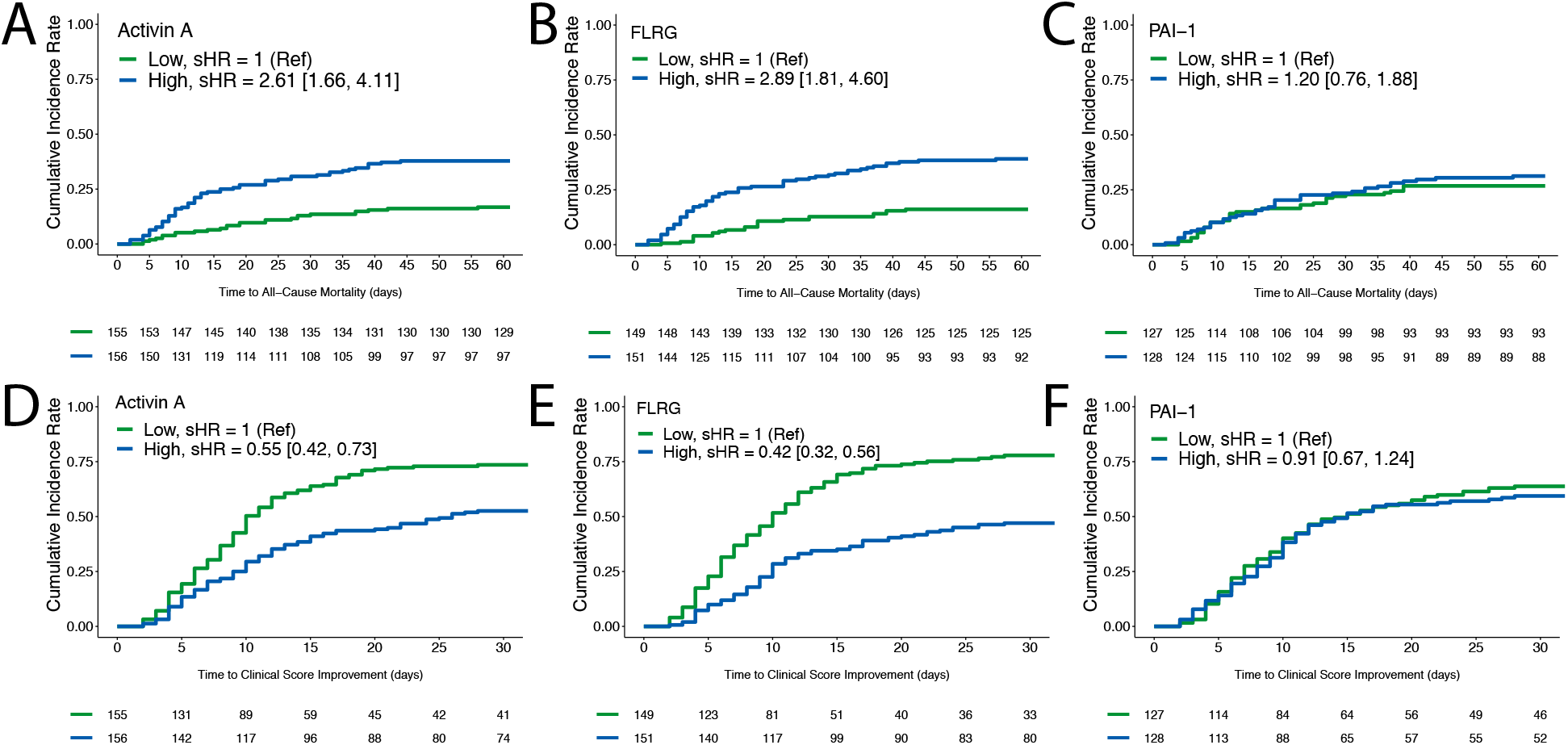
Cumulative incidence curves for all-cause mortality (A-C) and *≥*1-point clinical score improvement (D-F). All biomarkers (Activin A, FLRG, and PAI-1) were median split to form high and low groups. Low groups are in green and high groups are in blue. A-C: Cumulative incidence rate of all-cause mortality is plotted over time. D-F: Cumulative incidence rate of clinical score improvement (*≥*1 point) is plotted over time. Number of subjects remaining at risk for each event are shown at 5-day intervals starting at study enrollment.

**Table 3.** **Fine-Gray subdistribution hazard model results**.

|  | **Activin A** |  | **FLRG** |  | **PAI-1** |  |
| --- | --- | --- | --- | --- | --- | --- |
|  | **Low** | **High** | **Low** | **High** | **Low** | **High** |
| No. patients<sup>#</sup> | 156 | 156 | 150 | 151 | 128 | 128 |
| **All-cause mortality** |  |  |  |  |  |  |
| No. events | 26 (17%) | 59 (38%) | 24 (16%) | 59 (39%) | 34 (27%) | 40 (31%) |
| Person-days | 7973 | 6650 | 7754 | 6300 | 6044 | 5860 |
| Event rate* | 3.26<br>(2.18, 4.71) | 8.87<br>(6.82, 11.4) | 3.10<br>(2.03, 4.54) | 9.37<br>(7.19, 12.0) | 5.63<br>(3.96, 7.77) | 6.82<br>(4.94, 9.20) |
| Unadjusted sHR | 1 (Ref) | 2.61<br>(1.66, 4.11) | 1 (Ref) | 2.89<br>(1.81, 4.60) | 1 (Ref) | 1.20<br>(0.76, 1.88) |
| Adjusted sHR | 1 (Ref) | 2.82<br>(1.60, 4.95) | 1 (Ref) | 2.90<br>(1.50, 5.59) | 1 (Ref) | 1.08<br>(0.53, 1.64) |
| **Clinical score improvement** |  |  |  |  |  |  |
| No. events | 114 (73%) | 82 (53%) | 116 (77%) | 71 (47%) | 81 (63%) | 76 (59%) |
| Person-days | 2236 | 3067 | 2019 | 3114 | 2171 | 2247 |
| Event rate** | 5.10<br>(4.23, 6.10) | 2.67<br>(2.14, 3.30) | 5.75<br>(4.77, 6.87) | 2.28<br>(1.79, 2.86) | 3.73<br>(2.98, 4.61) | 3.38<br>(2.68, 4.21) |
| Unadjusted sHR | 1 (Ref) | 0.55<br>(0.42, 0.73) | 1 (Ref) | 0.42<br>(0.32, 0.56) | 1 (Ref) | 0.91<br>(0.67, 1.24) |
| Adjusted sHR | 1 (Ref) | 0.56<br>(0.42, 0.76) | 1 (Ref) | 0.46<br>(0.33, 0.63) | 1 (Ref) | 0.98<br>(0.69, 1.40) |

**Note:** The sample was median split for each analyte separately and subdistribution hazard ratios (sHR) along with 95% confidence intervals were calculated for High relative to Low groupings. sHR>1 indicates higher rates in subjects with elevated Activin A, FLRG, or PAI-1. sHR<1 indicates lower rates in subjects with elevated Activin A, FLRG, or PAI-1. The variables in the adjusted models include age, sex, race, ethnicity, steroid use, duration of pneumonia prior to baseline, BMI, diabetes, hypertension, and treatment arm as covariates. \* per 1000 person-days, \*\* per 100 person-days. <sup>#</sup> 1 subject with missing longitudinal data in Low Activin A, FLRG, and PAI-1 groups.

COVID-19 patients with elevated Activin A were more likely to die than patients with low Activin A (sHR = 2.61 (1.66, 4.11); *p* < 0.0001). A similar effect was observed for FLRG (sHR = 2.89 (1.81, 4.60); *p* < 0.0001). However, no differences in mortality were observed based on a median split of PAI-1 (sHR = 1.20 (0.76, 1.88); *p* = 0.44). At day 60 of the study, rates of mortality were 17% (Low) and 38% (High) for Activin A, 16% (Low) and 39% (High) for FLRG, and 27% (Low) and 31% (High) for PAI-1.

COVID-19 patients with elevated Activin A were less likely to achieve clinical score improvement (*≥*1 point) than patients with low Activin A (sHR = 0.55 (0.42, 0.73); *p* < 0.0001). A similar effect was observed for FLRG (sHR = 0.42 (0.32, 0.56); *p* < 0.0001). However, no differences in clinical improvement were observed based on a median split of PAI-1 (sHR = 0.91 (0.67, 1.24); *p* = 0.56). At day 29 of the study, rates of clinical score improvement (*≥*1 point) were 73% (Low) and 53% (High) for Activin A, 77% (Low) and 47% (High) for FLRG, and 63% (Low) and 59% (High) for PAI-1.

The clinical data established that Activin A and its pathway marker FLRG were elevated in the most severe settings of Covid-19 and that high levels were predictive of the worst Covid19 outcomes; we therefore were interested in further investigating the relationship between the cytokines elevated in the Covid19 “cytokine storm” and Activin A. We studied cell types of particular interest given the pathology of Covid19 - bronchial and pulmonary smooth muscle cells.

### IL-1*α* and TNF*α* induce Activin A & FLRG in human bronchial/tracheal smooth muscle cells, lung fibroblasts, and pulmonary artery smooth muscle cells

We next sought to investigate cells that are relevant for COVID-19 and decided upon studying bronchial/tracheal smooth muscle cells and pulmonary artery smooth muscle cells, to determine if these produced Activin A in response to inflammatory cytokines. These cells seemed especially relevant since the prior cited paper documented very high levels of Activin A in the bronchial alveolar lavage fluid of ARDS patients (Apostolou et al., 2012). A third cell line that we thought to test were bronchial fibroblasts. We included these because it has been shown that COVID-19 is more severe in the aged, and those with pre-existing conditions such as COPD (Fang et al., 2020; Wang et al., 2020), and fibroblasts are more prevalent in these settings. These cell types are vulnerable or culprit cell populations in numerous pulmonary diseases – for example, they’re contributors to pulmonary remodeling and functional decline in chonic inflammatory lung diseases. Thus human bronchial/tracheal smooth muscle cells (SMCs), lung fibroblasts and pulmonary artery SMCs were treated with 10ng/ml IL-1*α* and TNF*α* for five days (Fig. 4). We found that both IL-1*α* and TNF*α* stimulated Activin A production in bronchial/tracheal SMC, pulmonary artery SMCs and lung fibroblasts and IL-1*α* had a more potent effect on Activin A production in comparison to TNF*<u>α</u>* (Fig. 4B).

**Figure 4:**
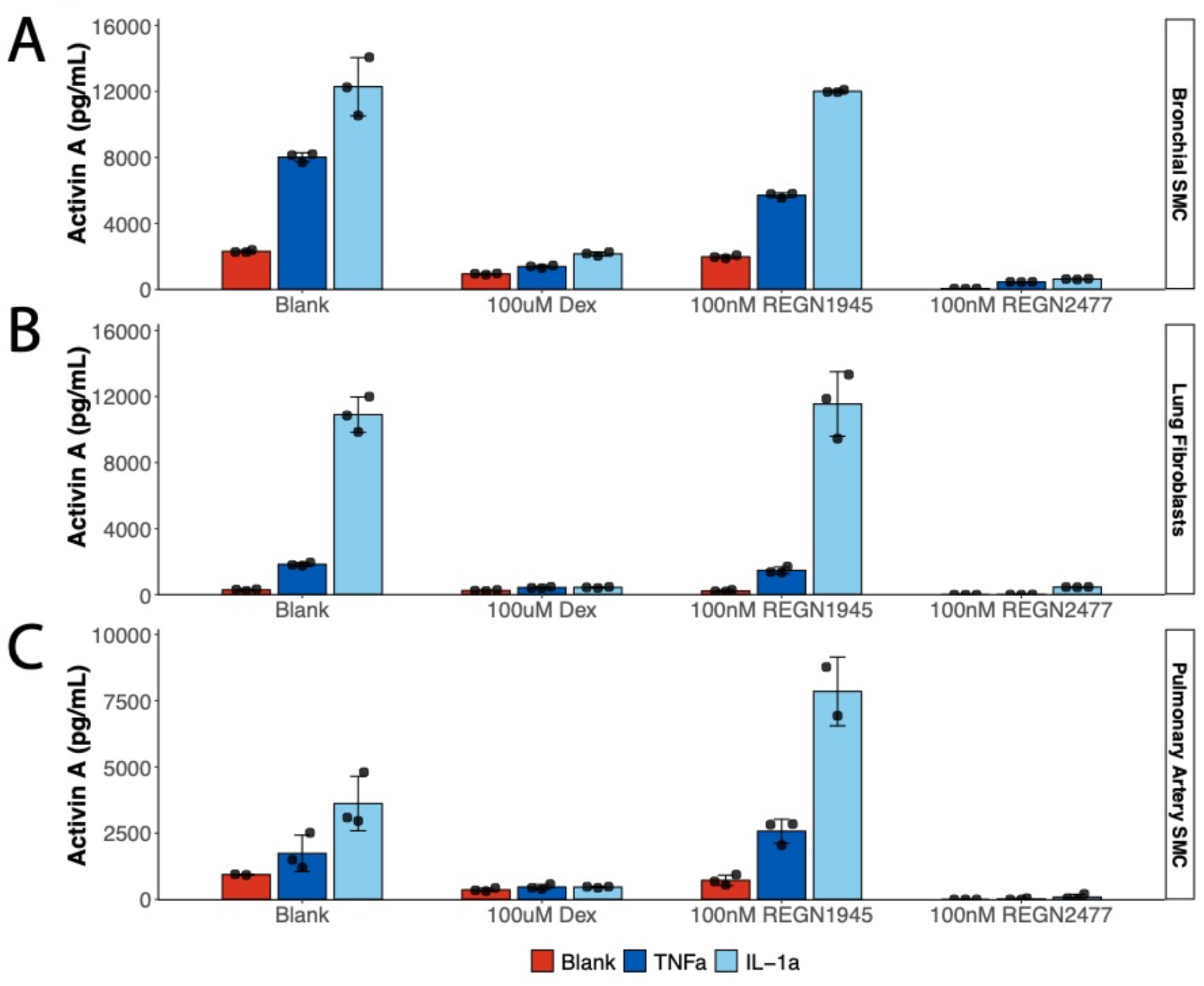
Both IL-1*α* and TNF*α* can stimulate Activin A production in bronchial/tracheal SMCs, lung fibroblasts and pulmonary artery SMCs, which can be rescued by dexamethasone. Activin A production was measured in conditioned medium by enzyme-linked immunosorbent assay (ELISA). BTSMC (Bronchial/tracheal smooth muscle) cells, NHLF (Normal human Lung Fibroblasts) and PASMC (Pulmonary artery smooth muscle cells) were serum starved overnight and then treated with 10ng/ml IL1 *α* or TNF*α* for five days in the presence or absence of 100uM Dexamethasone, 100nM anti-Activin A antibody REGN2477 or isotype control REGN1945. Conditioned medium was diluted at 1:10 for ELISA. The experiment was repeated twice, with a consistent result.

Dexamethasone, a corticosteroid used in wide range of conditions for its anti-inflammatory and immunosuppressant effect, has been tested in hospitalized COVID-19 patients and found to have benefits for critically ill patients. Here we also demonstrated that dexamethasone could reduce Activin A levels induced by IL-1*α* and TNF*α* back to baseline in bronchial/tracheal, pulmonary artery SMCs, and lung fibroblasts (Fig. 4). Although we used a higher dose (100uM) of dexamethasone in our *in vitro* study than can be used in the clinic, it is clear that dexamethasone at least is capable of inhibiting cytokine-induced induction of Activin A.

To see the effect of anti-Activin A, bronchial/tracheal and pulmonary artery SMC as well as lung fibroblasts were treated with an anti-Activin A ab (REGN2477) at 100nM, prior to the treatment with IL-1*α* or TNF*α*. Compared with the isotype control antibody (REGN1945), REGN2477 significantly reduced Activin A detection following treatment with IL-1*α* or TNF*α*, indicating that the antibody binds Activin A.

### IL-1*α* and TNF*α* induce Activin A via the IKK/NFkB pathway

We next wanted to further explore the mechanism by which inflammatory cytokines can induce Activin A, in pulmonary artery and in bronchial smooth muscle cells, which are both relevant for Covid19 pathology – since SARS-Cov2 prominently effects the lung and the blood vessels (Fig. 5). In particular, we wanted to dissect the role of signaling molecules IKK, p38 and JNK that are induced in response to cytokine receptor activation.

**Figure 5:**
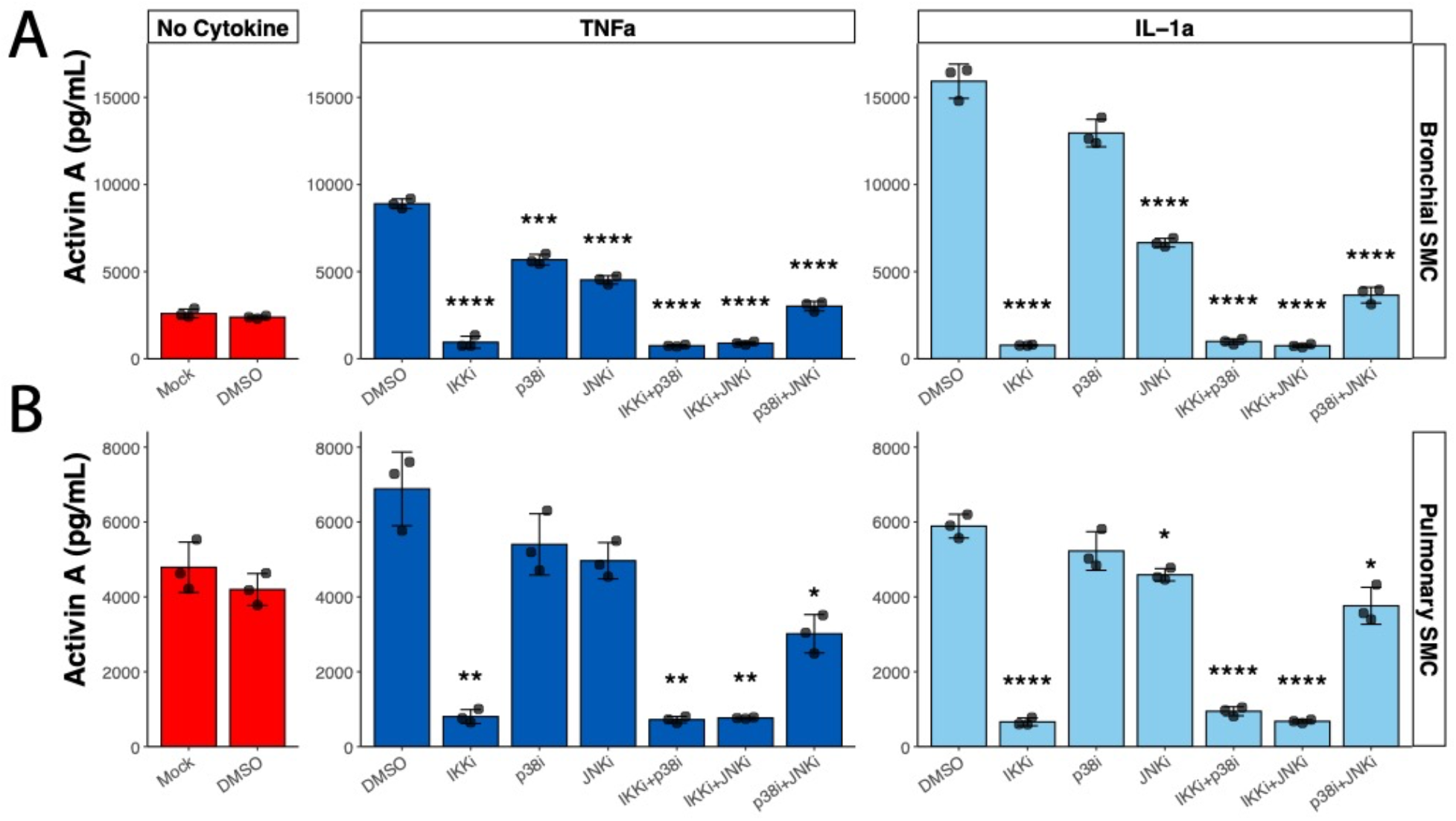
Role of IKK, p38 and JNK in Activin A induction in response to cytokine treatment. (A) Bronchial/tracheal smooth muscle cells (SMC) and (B) pulmonary artery smooth muscle cells (SMC) were treated for 24 hours with 100ng/ml of TNF*α* or IL1*α* in combination with pharmacological inhibitors for IKK (3uM Withaferin A), p38 (0.3uM SB203580) and JNK (30uM SP600125). A combination of IKK and p38 inhibitors (same concentrations), a combination of IKK and JNK inhibitors (same concentrations) and a combination of p38 and JNK inhibitors (same concentrations) was also used. DMSO was used as a negative control for inhibitor treatments. Treatment with sterile water only (Mock) was used as an additional control. Activin A levels were quantified in conditioned media by ELISA. N=3 for each treatment. Significant Bonferroni-corrected pairwise comparisons with DMSO within each treatment condition are labeled. * *p* < 0.05, ** *p* < 0.01, *** *p* < 0.001, **** *p* < 0.0001.

(Fig. In each cell type, the inflammatory cytokines could increase the production of Activin A, and this was blocked by inhibiting the IKK/NFkappaB pathway (Figs 5A and 5B). Inhibition of p38 or JNK could partially inhibit Activin A production, but this was cell-context specific, and not at all as complete as simply blocking IKK (Fig 5A-B). Furthermore, simultaneous inhibition of IKK and p38 or IKK and JNK was not additive, since IKK inhibition was sufficient to almost fully inhibit Activin A production (Figs. 5A-5B).

These results indicate that Activin A induction by IL1*α* and TNF*α* is mainly dependent on the IKK/NF-kappaB pathway and can be partially inhibited by blocking p38 or JNK pathways.

## Discussion

COVID-19 is a recent and potentially very serious disease; in the most severe settings it induces Acute Respiratory Disease Syndrome (ARDS), which can lead to the need for mechanical ventilation (Piroth et al., 2020). Severe cases of COVID-19 can also result in death; worldwide mortality from the disease ranges from 1.5 – 15%, depending on the country and other clinical risk factors. ARDS can be caused by a variety of other conditions, including sepsis (Confalonieri et al., 2017; Fialkow et al., 2006). In a prior survey of non-Covid19 ARDS patients, it was noted that there was a significant increase in levels of Activin A in the bronchial alveolar lavage fluid relative to control levels (Apostolou et al., 2012); these same researchers demonstrated that Activin A was sufficient to induce an ARDS-like phenotype in a pre-clinical model, by delivering the gene intra-tracheally via an adenovirus (Apostolou et al., 2012). A follow-up to this study was able to reproduce the prior finding, and demonstrated that Activin A and its pathway marker, FLRG (follistatin-like related gene, also called FLSTL3), were demonstrably elevated in the sera of non-Covid-19 ARDS patients (Kim et al., 2019). Such studies are clearly needed, to establish reliable biomarkers the reproducibility of biomarkers for ARDS; what is especially needed are biomarkers which predict disease severity so that clinicians can be alerted to the need for stronger interventions.

When patients with severe ARDS from COVID-19 began to be studied it was reported early on that these patients were experiencing a “cytokine storm”, which included reports of many inflammatory cytokines, including IL-1 and TNF*α* upregulated in the blood. Our group had previously connected IL-1 and TNF*α* to the induction of Activin A, in a study of skeletal muscle atrophy caused by these cytokines (Trendelenburg et al., 2012). We were interested in the mechanism of inflammation-induced atrophy, since inflammatory cytokines are upregulated in several settings of cachexia, including cancer cachexia, and the resultant loss of muscle is a serious contributor to mortality (Fearon et al., 2012). In our prior study we found that blocking Activin A could inhibit approximately 50% of the cytokine-induced muscle atrophy, demonstrating also in that setting that Activin A had a causal role for the downstream effects of inflammatory cytokines (Trendelenburg et al., 2012). In the lung ARDS study, it was also shown that inhibiting the Activin A pathway with a receptor trap could block the ability of LPS to induce ARDS (Apostolou et al., 2012), further strengthening the evidence that Activin A was required to induce ARDS, at least in a preclinical model.

We therefore sought in the present study to determine if Activin A and its pathway marker, FLRG, were upregulated in patients with COVID-19, and especially those who had ARDS, in comparison to age-matched controls. In this study we found that both Activin A and FLRG were upregulated quite significantly in ICU-bound Covid-19 patients - with some patients experiencing more than 2-fold increases in Activin A, with a tail of subjects that experienced increases as high as 8x above the normal median. Strikingly, Activin A levels were not significantly increased in COVID-19 patients who had serious disease but did not require invasive mechanical ventilation - the marker clearly distinguished the populations. Activin A induces FLRG (follistatin-related gene, also known as follistatin-like 3) (Bartholin et al., 2001). In contrast to Activin A, which is internalized - and thus may not be found in the blood at lower levels of pathway activation - FLRG remains in the blood stream, where it inhibits Activin function - thus it’s a downstream marker of prior activation of the pathway. FLRG levels were induced in the severe, non-ICU patients, but were even more elevated in the ICU patients, especially those who were experiencing ARDS, as determined by their need for invasive mechanical ventilation. Additionally, with FLRG, subjects with very high elevations were found to be more likely to require invasive mechanical ventilation as well.

The action of Activin and FLRG is likely not to be independent, with a correlation of (Spearman rho = 0.54, *p* < 0.0001) at baseline between the two analytes, providing statistical evidence that they’re likely within the same pathway, which has been demonstrated biologically (Bartholin et al., 2001). In contrast, PAI-1, which is a marker of coagulation, is already elevated even in less severely affected patients, and was not further elevated in the ICU patients. Therefore, the data in this paper establishes that Activin A and FLRG levels distinguish those who go on to the most serious form of the disease, as determined by the need for oxygen, and risk of dying from COVID-19. In contrast, PAI-1 is a less reliable biomarker for the eventual need for oxygenation, or the risk of mortality. Although the data does not itself provide evidence that elevated Activin/FLRG are causative of morbidly and mortality, it does indicate that these are appropriate targets for further analyses, and prior preclinical data was suggestive that Activin A may be causative for ARDS (Apostolou et al., 2012). In the present analysis, baseline levels of Activin A and FLRG (not PAI-1) predicted all-cause mortality both with and without inclusion of covariates in the models. A one standard deviation difference in Activin A (+374.1 pg/mL) at baseline resulted in a 54% increased likelihood of death and a one standard deviation difference in FLRG (+9415 pg/mL) resulted in a 69% increased likelihood of death. In a complementary analysis, patients with Activin A greater than the sample median (388.4 pg/mL) were 2.6 times more likely to die than patients with lower levels. Patients with FLRG greater than the sample median (13531 pg/mL) were 2.9 times more likely to die than patients with lower levels. These effects remained consistent when treatment arm and other covariates were included in the models.

Prior studies have shown that Activin A can cause fibrosis (Matsuse et al., 1996), which is seen in ARDS lungs, perhaps this is one way in which Activin A contributes to the phenotype. Further, activation of myostatin, a related ligand, was shown to block skeletal muscle differentiation (Trendelenburg et al., 2009); however these TGFbeta superfamily molecules can induce fibroblast proliferation (Ohga et al., 1996) – which is also suggestive of their possible role in this clinical setting. In addition, a paper looking at age-associated changes in protein levels in human plasma was recently published (Lehallier et al., 2019); when we analyzed Activin A (InhbA.InhbB) and FLRG (FSTL3) using their publicly available data (https://twc-stanford.shinyapps.io/aging_plasma_proteome/), both were shown to be significantly upregulated with age - this upregulation could lower the threshold for pathological upregulation induced by inflammatory cytokines.

We were interested to explore the mechanism by which inflammatory cytokines induce Activin A, and what the likely cell types were which might be responsible. Examination of cell lines demonstrated that IL-1 could induce Activin A significantly in Bronchial Smooth muscle cells, as well as pulmonary muscle cells. It also had this effect on lung fibroblasts, which might be more prevalent in the aged, or those with pre-existing conditions, such as COPD. TNF*α* was a less reliable inducer of Activin A infibroblast cell lines; we speculate that this may be due to relative levels of IL-1 vs TNF receptors, rather than some intrinsic difference in the signaling pathways - we note this because TNF*α* is fully capable of inducing Activin A in pulmonary and bronchial smooth muscle cells - so there doesn’t seem to be anything intrinsically different between the two cytokines, just a cell-specific ability to respond. As to downstream mechanisms responsible for the induction of Activin A, we distinguished between various signaling pathway induced by IL-1 and TNF using cellular inhibitors of those pathways. Only the IKK inhibitor was able to completely and consistenly block the induction of Activin A, as opposed to p38 or Jnk inhibitors, which were less effective. IKK is upstream of NF-kappaB induction, so the data demonstrates that Activin A is induced by this pathway.

The cellular differences in response to IL-1 vs TNF demonstrate that a cytokine-centric approach is unlikely to be successful to treat a disease like COVID-19, where there are a large array of cytokines induced. Many of these induced cytokines signal similarly, and therefore one cannot hope to block the ultimate effect without going downstream in the signaling pathway. Thus, it is of particular interest that Activin A is induced by the IKK/NF-kappaB pathway, in common to IL-1 and TNF, and other inflammatory cytokines, and that Activin A is sufficient, at least in a pre-clinical model to induce ARDS. Furthermore, since Activin A and its pathway marker, FLRG, are unique in being associated with the most severe effects caused by COVID-19 (the induction of ARDS, resulting in an inability to breathe, ultimately causing death), the strong suggestion from this data is that it would be beneficial to COVID-19 patients suffering from ARDS to be treated with an inhibitor of Activin A.

Of note, an antibody to Activin A was fully capable of blocking the measured Activin A levels induced by inflammatory cytokines, giving further hope that such an antibody could be effective in blocking the severe effects of COVID-19 in ICU patients. Indeed, in the previously published preclinical setting, use of an Activin receptor trap was sufficient to ameliorate ARDS in an LPS-induced model of the syndrome (Apostolou et al., 2012).

Might there be negative consequences to using an anti-Activin A approach in ARDS patients? One study did show that Activin A could inhibit viral production in a variety of virus-infected cell lines (Eddowes et al., 2019). This seems to be a plausible evolutionary mechanism explaining why Activin A might be induced by cytokines. Initial viral load does correlate with disease severity (Fajnzylber et al., 2020). However, the cytokine storm often comes later in the disease, indicating the cytokine storm is part of the immune response to the virus (Khadke et al., 2020). The impression, therefore, is that these severely affected patients are not suffering directly from the viral load, but instead from an over-reaction of the immune system - the later occurring cytokine response, which in some patients over-induces Activin A. We therefore suggest it’s reasonable to try inhibiting Activin A or its induced pathway to treat COVID-19 patients who are experiencing ARDS.

## Acknowledgments

We thank Drs. G.D. Yancopoulos, A. Murphy, L. S. Schleifer and P. Roy Vagelos for enthusiastic support, along with the rest of the Regeneron community, particularly the Sarilumab 2040 study team and site staff and patients who participated in the study. We thank Alpana Waldron and Rafia Bhore for assistance with the clinical database, Jennifer Hamilton for advice, Scott Macdonnell for sharing data and giving advice, and Kishor Devalaraja, for discussions. We’re particularly indebted to the rest of the Aging/Age-Related Disorders Group for valuable discussions. Thank you also to D. Lederer for critically reading the manuscript. Certain aspects of this project have been funded in whole or in part with Federal funds from the Department of Health and Human Services; Office of the Assistant Secretary for Preparedness and Response; Biomedical Advanced Research and Development Authority, under OT number: HHSO100201700020C.

## Author contributions

Megan McAleavy and Qian Zhang: design of experiments, ELISAs, cell experiments; manuscript preparation.

Jianing Xu and Matthew Wakai: mechanistic experiments (NF-kappaB pathway); manuscript preparation.

Li Pan: design and organization of experiments.

Peter J. Ehmann, Matthew F. Wipperman, Sara C. Hamon, Anita Boyapati: Statistical analyses and manuscript preparation.

Tea Shavlakadze: design of experiments and manuscript preparation.

Lori G. Morton, Christos A. Kyratsous: design of experiments.

David J. Glass: project conception; design of experiments and manuscript preparation.

## Methods

### COVID-19 Samples and informed consent

Samples were collected from subjects who consented to participate in an adaptive, phase 2/3, randomized, double-blind, placebo-controlled trial of intravenous (IV) sarilumab in adults hospitalized with severe or critical COVID-19. In the phase 3 portion, patients with critical COVID-19 were randomized to sarilumab 400 mg IV, sarilumab 200 mg IV, or placebo (Trial registration number: NCT04315298). The protocol was developed by the sponsor (Regeneron Pharmaceuticals Inc.). Data were collected by the study site investigators and analyzed by the sponsors. The local institutional review board or ethics committee at each study center oversaw trial conduct and documentation. All patients provided written informed consent before participating in the trial.

### Definition of clinical variables

**Oxygen Device Type:** High Flow O2 requirements include: Non-rebreather face mask, High-flow nasal cannula, Non-invasive ventilation. Low Flow O2 requirements include Nasal cannula, simple face mask. **All-cause mortality:** Number of days from randomization to death. **Clinical Improvement:** ≥1-point improvement in clinical status from baseline to Day 22 using the Clinical Status Assessment (7-point ordinal scale) (Peterson, 2017): 1, death; 2, hospitalized, requiring invasive mechanical ventilation membrane; 3, hospitalized, requiring non-invasive ventilation or high-flow oxygen devices; 4, hospitalized, requiring supplemental oxygen; 5, hospitalized, not requiring supplemental oxygen but requiring ongoing medical care (COVID-19 related or otherwise); 6, hospitalized, not requiring supplemental oxygen and no longer requiring ongoing medical care; and 7, not hospitalized.

### Reagents

Withaferin A (Cat # S8587), SB203580 (Cat # S1076), SP600125 (Cat # S1460) were acquired from Selleckchem. Recombinant Human TNF*α* (Cat # 300-01A), IL1b (Cat # AF-200-01B), IL1a (Cat #200-01A) were acquired from Peprotech.

### Cell Culture

#### Cytokine and steroid treatments

Bronchial/Tracheal smooth muscle cell (BTSMC, ATCC, PCS-130-011), normal human lung fibroblast (NHLF, Lonza, CC-2512), and pulmonary artery smooth muscle cell (PASMC, Lonza, CC-2581) were seeded at 200,000 cells/well in twelve-well plates and cultured in media as specified by manufacturers. To eliminate the influence of other factors, cells were washed once and cultured with serum-free medium with 5% BSA overnight. To study the effect of pro-inflammatory cytokines IL-1*α* and TNF*α* on Activin A production and whether Dexamethasone and anti-Activin A can rescue, cells were pretreated with 100µM dexamethasone (Sigma-Aldrich Inc, #D2915), 100nM REGN1945 (isotype control) or 100nM REGN2477 (anti-Activin A) for 10 minutes, and then treated with or without 10ng/ml IL-1*α* (R&D Systems, #200-LA-CF) or TNF*α* (R&D Systems, #210-TA-CF) for five days. Conditioned media were harvested and diluted at 1:10, and ELISA (human/mouse/rat activin A Quantikine ELISA kit, R&D Systems, #DAC00B) was performed to measure Activin A production according to the manufacturer’s protocol.

#### ELISAs

Activin A, FLRG, and PAI-1 ELISAs were carried out by using R&D Quantikine ELISA Kit systems, DAC00B, DFLRG0, DSE100 respectively, according to manufacturer’s protocols.

#### Statistical analysis

*Clinical study:* Descriptive statistics grouped by disease severity and oxygenation requirements are reported as median (interquartile range; IQR) for continuous variables and frequency (percent; %) for categorical variables. Kruskal Wallis tests were used for continuous variables across three or more groups and Wilcoxon signed-rank tests were used for continuous variables between two groups. Fisher’s exact tests were used for categorical variables. Kruskal Wallis tests with follow-up Dunn pairwise comparisons (for significant omnibus results) were used to test for differences in Activin A, FLRG, and PAI-1 by disease severity and oxygen requirements at baseline. For longitudinal outcomes, including mortality and clinical score improvement (*≥*1 point), logistic regression models were used with baseline Activin A, FLRG, and PAI-1 (standardized) as predictors, with and without inclusion of covariates. Fine-Gray subdistribution hazard models were also generated for these longitudinal outcome variables with available time-to-event information. Subjects were split into Viral Load groupings (Low, High) based on a median split of baseline values of each analyte individually. Subdistribution hazard ratios (sHR) for High groups, relative to Low groups, were calculated with and without inclusion of covariates. All-cause mortality and clinical score improvement time-to-event data were censored to Day 60 and Day 29, respectively. Rates of incidence for each outcome during the study were calculated at the censored timepoint.

##### Experiment 1

Within each treatment condition (Blank, TNF*α*, and IL-1*α*) and cell type (Bronchial SMC, Lung Fibroblasts, Pulmonary Artery SMC), pairwise *t*-tests were used to compare Activin A concentration after additional cotreatment of 100uM Dex, 100nM REGN, and 100nm REGN2477 to no cotreatment (Blank).

##### Experiment 2

Within each main treatment condition (no cytokines, IL1*α*, TNF*α*) and cell type (bronchial smooth muscle, pulmonary smooth muscle), Activin A induction for each cotreatment was compared to DMSO cotreatment using pairwise *t*-tests.

A Type-I error rate of *α* = 0.05 was used as the threshold for statistical significance, with Bonferroni adjustment for follow-up tests and multiple comparisons. Covariates for all indicated analyses included age, sex, race, ethnicity, steroid use, duration of pneumonia prior to baseline, BMI, diabetes, and hypertension. Treatment arm was included as a covariate in models predicting longitudinal outcomes. The results were almost identical regardless of inclusion or exclusion of covariates; therefore, unadjusted statistics are reported in the text. All statistical analyses were performed using R version 3.6.1.

## References

Ahmed, S., Zimba, O., and Gasparyan, A.Y. (2020). Thrombosis in Coronavirus disease 2019 (COVID-19) through the prism of Virchow’s triad. Clin Rheumatol 39, 2529–2543.

Apostolou, E., Stavropoulos, A., Sountoulidis, A., Xirakia, C., Giaglis, S., Protopapadakis, E., Ritis, K., Mentzelopoulos, S., Pasternack, A., Foster, M., et al. (2012). Activin-A Overexpression in the Murine Lung Causes Pathology That Simulates Acute Respiratory Distress Syndrome. American Journal of Respiratory and Critical Care Medicine 185, 382–391.

Bartholin, L., Maguer-Satta, V., Hayette, S., Martel, S., Gadoux, M., Bertrand, S., Corbo, L., Lamadon, C., Morera, A.-M., Magaud, J.-P., et al. (2001). FLRG, an activin-binding protein, is a new target of TGFβ transcription activation through Smad proteins. Oncogene 20, 5409–5419.

Bos, L.D., Schouten, L.R., van Vught, L.A., Wiewel, M.A., Ong, D.S.Y., Cremer, O., Artigas, A., Martin-Loeches, I., Hoogendijk, A.J., van der Poll, T., et al. (2017). Identification and validation of distinct biological phenotypes in patients with acute respiratory distress syndrome by cluster analysis. Thorax 72, 876–883.

Calfee, C.S., Janz, D.R., Bernard, G.R., May, A.K., Kangelaris, K.N., Matthay, M.A., and Ware, L.B. (2015). Distinct molecular phenotypes of direct vs indirect ARDS in single-center and multicenter studies. Chest 147, 1539–1548.

Confalonieri, M., Salton, F., and Fabiano, F. (2017). Acute respiratory distress syndrome. European Respiratory Review 26, 160116.

Eddowes, L.A., Al-Hourani, K., Ramamurthy, N., Frankish, J., Baddock, H.T., Sandor, C., Ryan, J.D., Fusco, D.N., Arezes, J., Giannoulatou, E., et al. (2019). Antiviral activity of bone morphogenetic proteins and activins. Nat Microbiol 4, 339–351.

Egerman, M.A., and Glass, D.J. (2014). Signaling pathways controlling skeletal muscle mass. Critical Reviews in Biochemistry and Molecular Biology 49, 59–68.

Fajnzylber, J., Regan, J., Coxen, K., Corry, H., Wong, C., Rosenthal, A., Worrall, D., Giguel, F., Piechocka-Trocha, A., Atyeo, C., et al. (2020). SARS-CoV-2 viral load is associated with increased disease severity and mortality. Nature Communications 11, 5493.

Fang, X., Li, S., Yu, H., Wang, P., Zhang, Y., Chen, Z., Li, Y., Cheng, L., Li, W., Jia, H., et al. (2020). Epidemiological, comorbidity factors with severity and prognosis of COVID-19: a systematic review and meta-analysis. Aging 12, 12493–12503.

Fearon, K. C H., Glass, D., and Guttridge, D. (2012). Cancer Cachexia: Mediators, Signaling, and Metabolic Pathways. Cell Metabolism 16, 153–166.

Fialkow, L., Fochesatto Filho, L., Bozzetti, M.C., Milani, A.R., Rodrigues Filho, E.M., Ladniuk, R.M., Pierozan, P., de Moura, R.M., Prolla, J.C., Vachon, E., et al. (2006). Neutrophil apoptosis: a marker of disease severity in sepsis and sepsis-induced acute respiratory distress syndrome. Crit Care 10, R155–R155.

Huang, C., Wang, Y., Li, X., Ren, L., Zhao, J., Hu, Y., Zhang, L., Fan, G., Xu, J., Gu, X., et al. (2020). Clinical features of patients infected with 2019 novel coronavirus in Wuhan, China. Lancet (London, England) 395, 497–506.

Jaynes, J.B., Johnson, J.E., Buskin, J.N., Gartside, C.L., and Hauschka, S.D. (1988). The muscle creatine kinase gene is regulated by multiple upstream elements, including a muscle-specific enhancer. Mol Cell Biol 8, 62–70.

Khadke, S., Ahmed, N., Ahmed, N., Ratts, R., Raju, S., Gallogly, M., de Lima, M., and Sohail, M.R. (2020). Harnessing the immune system to overcome cytokine storm and reduce viral load in COVID-19: a review of the phases of illness and therapeutic agents. Virology Journal 17, 154.

Kim, J.M., Lee, J.K., Choi, S.M., Lee, J., Park, Y.S., Lee, C.H., Yim, J.J., Yoo, C.G., Kim, Y.W., Han, S.K., et al. (2019). Diagnostic and prognostic values of serum activin-a levels in patients with acute respiratory distress syndrome.

Lehallier, B., Gate, D., Schaum, N., Nanasi, T., Lee, S.E., Yousef, H., Moran Losada, P., Berdnik, D., Keller, A., Verghese, J., et al. (2019). Undulating changes in human plasma proteome profiles across the lifespan. Nature Medicine 25, 1843–1850.

Matsuse, T., Ikegami, A., Ohga, E., Hosoi, T., Oka, T., Kida, K., Fukayama, M., Inoue, S., Nagase, T., Ouchi, Y., et al. (1996). Expression of immunoreactive activin A protein in remodeling lesions associated with interstitial pulmonary fibrosis. The American journal of pathology 148, 707–713.

Ohga, E., Matsuse, T., Teramoto, S., Katayama, H., Nagase, T., Fukuchi, Y., and Ouchi, Y. (1996). Effects of Activin A on Proliferation and Differentiation of Human Lung Fibroblasts. Biochemical and Biophysical Research Communications 228, 391–396.

Piroth, L., Cottenet, J., Mariet, A.-S., Bonniaud, P., Blot, M., Tubert-Bitter, P., and Quantin, C. (2020). Comparison of the characteristics, morbidity, and mortality of COVID-19 and seasonal influenza: a nationwide, population-based retrospective cohort study. The Lancet Respiratory Medicine.

Portier, G.L., Benders, A.G., Oosterhof, A., Veerkamp, J.H., and van Kuppevelt, T.H. (1999). Differentiation markers of mouse C2C12 and rat L6 myogenic cell lines and the effect of the differentiation medium. In Vitro Cell Dev Biol Anim 35, 219–227.

Razanajaona, D., Joguet, S., Ay, A.-S., Treilleux, I., Goddard-Léon, S., Bartholin, L., and Rimokh, R. (2007). Silencing of FLRG, an Antagonist of Activin, Inhibits Human Breast Tumor Cell Growth. Cancer Research 67, 7223.

Trendelenburg, A.U., Meyer, A., Rohner, D., Boyle, J., Hatakeyama, S., and Glass, D.J. (2009). Myostatin reduces Akt/TORC1/p70S6K signaling, inhibiting myoblast differentiation and myotube size. American Journal of Physiology-Cell Physiology 296, C1258–C1270.

Trendelenburg, A.U., Meyer A Fau - Jacobi, C., Jacobi C Fau - Feige, J.N., Feige Jn Fau - Glass, D.J., and Glass, D.J. (2012). TAK-1/p38/nNFκB signaling inhibits myoblast differentiation by increasing levels of Activin A.

Wang, B., Li, R., Lu, Z., and Huang, Y. (2020). Does comorbidity increase the risk of patients with COVID-19: evidence from meta-analysis. Aging 12, 6049–6057.

